# Essential oil composition of *Callistemon citrinus* (Curtis) and its protective assessment towards *Tribolium castaneum* (Herbst) (Coleoptera: Tenebrionidae)

**DOI:** 10.1101/2021.01.05.425383

**Authors:** Maduraiveeran Ramachandran, Kathirvelu Baskar, Abeer Hashem, Elsayed Fathi Abd_Allah, Manickkam Jayakumar

## Abstract

Essential oil (EO) was extracted from *Callistemon citrinus* leaves by hydro-distillation. The extracted oil was analysed by GC and Mass Spectroscopy. Analysis report showed that the major constituent of the essential oil was eucalyptol (40.44%). The EO of *C. citrinus* exhibited 100% fumigation toxicity (adult mortality) against adult and 95.8% larvicidal activity against *Tribolium castaneum* at 160 μL/L (12 hrs) and 320 μL/L (48 hrs), respectively. The effective concentration of 37.05 μL/L (adult) and 144.31 μL/L (larva) at 24 and 48 hrs respectively. A 100% repellent activity was observed at 20 μl for adult beetles and 93.3% for larvae of *T. castaneum* at 20 μl after 24 h. Exposure to *C. citrinus* EO significantly reduced beetle fecundity, ovicidal activity, egg hatching, larvae survival, and emergence of adult. The effect of EO on detoxification enzymes of *T. castaneum* adults was examined. Results indicated that the activity of detoxification enzymes drastically varied when compared with control. This EO had toxicant effects on all stages of the life of *T. castaneum*. Hence it may be used as fumigant instead of the use of using synthetic chemical fumigants.

## Introduction

Insect infestation on stored grains, pulses, and their processed products is a major problem that results in significant economic losses and reduces the quality as well as the quantity of stored food products. Stored grains can be infested by several insect pests that cause severe damage. Storage pests alone damaged 14-17 million tonnes of food grains and nearly 15 insect species have been are listed as major stored grain pests in India [1]. Amongst, *Tribolium castaneum*, listed in the major pest category of stored grains, although predominantly found in tropical countries. Both larvae and adult of *T. castaneum* feed grains, seeds, and milled commodities. This beetle is responsible for approximately 10-40% of post-harvest losses worldwide [2], while in India estimates of losses range from 7-10% [3].

Fumigation is an effective method of pest management in stored grains. This method is used to control all stages of insects in stored grains and is cost effective, rapidly killing insects and leaving residues [4]. Currently, methyl bromide (CH_3_Br) and alumium phosphide (AlP) are approved for use on stored grain and are used as synthetic fumigants. However, Methyl bromides, has ozone depleting properties [5–7]. and insect resistance to phosphine has been documented [8–9]. Application of fumigants leads to increasing pest resurgence, deleterious effects on beneficial organisms, as well as raising the levels of toxicity [10]. To address these problems naturally biodegradable plant products have been evaluated.

Interestingly, plant leaves were used as a stored grain protectant in ancient times. Traditionally, plant-derived oils were used to protect stored pulses. More recently, essential oils (EOs) derived from plant have been receiving more attention as an alternatives to synthetic fumigants. Plant products protect food grains through their insecticidal and repellent properties. Plant-derived products are also generally harmless to flora and fauna in the environment. Thus, many researchers tested the plant essential oils for their biological potential against pests of stored food grains [11–14]. Essential oils have been extracted from members of the Myrtaceae, Lauraceae, Umbelliferae, Lamiaceae, Asteraceae, and from conifers [15–16].

Essential oils can exhibit fumigation toxicity, repellent activity, pupicidal activity, ovicidal, and oviposition deterrents against insect pests of stored grains [17–22]. Hence the present investigation was aimed to evaluate toxicity effect of *Callistemon citrinus* essential oil against *Tribolium castaneum*.

## Materials and Methods

### Culture of insect

The *Tribolium castaneum,* was maintained in Insectarium, Department of Zoology, University of Madras, and cultured on wheat flour. Freshly laid eggs, emerged larvae, and adults were used in the experiments.

### Chemicals

The chemical used in the study were analytical grade and were purchased from Sigma-Aldrich and Sisco Research Laboratories Pvt. Ltd. (India).

### Oil Extraction and GC-MS analysis

*Callistemon citrinus* (Bottle brush) was collected from the campus of University of Madras. The oil was extracted from freshly-collected plant leaves by hydro-distillation, then it was subjected to GC-MS analysis.

### Toxicity Study

Fumigation toxicity of *C. citrinus* EO was tested in the laboratory using filter paper method on adult insect at 28 ± 2°C and 60–70% RH [23]. Two different ranges of concentrations such as 40, 80, 120, 160 and 200 μl/L, and 40, 80, 120, 160, 200, 240, 280, and 320 μl/L air were evaluated for fumigation toxicity on adults and larvae, respectively. Ten freshly-emerged 3-7-day-old adult/10-12 days, old larvae were released in a bottle along with a small amount of flour as feed. The EO was poured on filter paper and it was adhered inside the screw cap of the bottle then closed tightly. Without treatment of essential oil to be consider control. Five replications were made for all treatments and controls. Mortality of adults and larvae were recorded after 3, 6, 9, 12, 24, 36, and 48 h commencement of treatments. Dead insects were counted, if there were no antennal or leg movements. Mortality was calculated using the Abbott formula [24].

### Repellency - Larvae

The repellency of EO was measured using the diet impregnation method on larvae. Twenty-five larvae per Petri dish and replicated five times. After 2, 4, 6, 12 and 24 h of treatment the number of larvae present in the treated and control diets were counted.

### Repellency - adult

The repellency effect of EO was evaluated with the help of glass olfactometer (Y- tube) on adult insect. Two grams of medium was mixed with different concentration of 5, 10, 20, and 30 μl of EO individually in each vial and attached into an arm of the olfactometer. Medium without essential oil was used as a control. All the glass vials attached to the arms and then fifty freshly-emerged adults were released into the olfactometer via the central opening. The number of beetles found in each vial was recorded after 24 hrs [25]. The percentage of repellent activity was calculated [26].

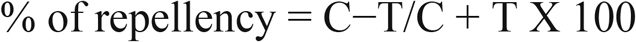

C- control; T- treatment

### Fecundity and knock-down effect

Ten adults were released into Petri dish (100 ml) with known quantity of wheat flour. Filter paper (Whatman No. 1) discs measuring about 2 cm dia, were impregnated with different concentrations (5, 10, 20 and 30 μL/L) of *C. citrinus* EO. At 24 hrs after treatment the adult were transferred to new Petri dish with food. The Petri dishes were carefully examined and recorded the number of eggs laid in control and treatments for a period of 2 days by using compound microscope. Knock-down adults were counted separately and recorded as a knock-down effect. Five replications were used for each treatment and the control group.

### Growth inhibition effects

Fifty adult beetles released in a Petri dish containing known quantity of wheat flour. After 48 h, the adult beetles were removed and numbers of eggs laid in each Petri dish were counted. Subsequently, the filter paper treated with different sub-lethal concentrations of EO was placed inside a Petri dish. Filter paper discs devoid of any volatiles were used as a control. The experiments were replicated five times for both treatments and control. The eggs hatched in each Petri dish was recorded daily and were maintained continuously on wheat flour. The larval survivability & per cent adult emergence (F_1_) were recorded.

### Sample preparation for biochemical studies

Adult insects were treated with sub-lethal concentrations of 5, 10 and 20 μL/L of EO. The live insects were used in the biochemical analysis which consisted of three replicates. Treated adults (10 individuals for each concentration) were transferred separately and homogenized with 500 μl of ice-cold phosphate buffer (20 mM, pH 7.0) using a Teflon, hand homogenizer to estimate the total protein, esterase, phosphatase, and Glutathione-S-Transferase activity. The homogenates were centrifuged at 15,000 rpm at 4 °C for 20 min. and the clear supernatants were stored at −20 °C until used. The supernatants were used for both qualitative and quantitative analyses.

### Biochemical analyses

Biochemical studies were carried out using previously described methods. The Bradford assay was used to determine total protein [27], acetylcholinesterase activity [28–29], α and β carboxylesterase [30] activity was estimated [31], levels of acid phosphatases (ACP) and alkaline phosphatises [32] were determined using the method of Koodalingam et al. [31] and Glutathione-S-Transferase activity was measured by Brogden and Barber [33] method.

### Estimation of biochemical components

The total protein, acid & alkaline phosphatases and ß-carboxylesterase of adult were examined by discontinuous PAGE gel using non-denatured conditions. The gel electrophoresis was run by using 5% stacking gel (pH 6.8) and 8% separating gel (pH 8.8) in Tris-glycine buffer (pH 8.3). The page was provided constant current of 4 mA per sample at 10°C on a slab gel; then it was stained.

### Estimation of esterase and phosphates activity

The α & β-carboxylesterase activity were detected by using separated protein bands by the method of Kirkeby and Moe [34]; Argentine and James [35]. Acid and alkaline phosphatase activitied were analysed as describe by Houk and Hardy [36].

### Statistical analysis

Student’s ‘t’ test was carried out to determine the significant differences between the biochemical constituents and enzyme activity in the treatments and control. Differences between means were considered as significant at p ≤ 0.05. All statistical analyses done original data (after transformed also the data did not showed significant distribution Shapiro wilk test). The probit analysis was done for fumigation toxicity. Significant different between the treatment group was calculated Duncans test followed by F-Test. The SPSS software, version 25was used for analysis.

## Results

### Oil yield

EO of *C. citrinus* was extracted from the leaves using a Clevenger apparatus at 65 °C for 3 h. Initially the oil was whitish in colour but later turned a pale yellow. The maximum yield of 650 μL/100 g (fresh weight of leaves) was obtained.

### Chemical composition of essential oils

Chemical composition of the *C. citrinus* EO was analysed by GC-MS and identified 10 different compounds in varying quantities. Among the 10 compounds, eucalyptol represented the major constituent (40.44%), followed by linalool (27.35%), and alfa- Pinene (17.36%) (Table 1).

**Table 1.** Phytochemical Constituents and composition (%) of essential oil from *Callistemon citrinus*.

| Peak No. | Compounds | Retention Time (min) | Area% |
| --- | --- | --- | --- |
| 1 | alpha.-Pinene | 5.690 | 17.46 |
| 2 | Camphene | 5.955 | 0.59 |
| 3 | beta.-Pinene | 6.412 | 5.83 |
| 4 | beta.-Myrcene | 6.627 | 3.22 |
| 5 | beta.-Pinene | 6.852 | 0.93 |
| 6 | Eucalyptol | 7.315 | 40.44 |
| 7 | gamma.-Terpinene | 7.738 | 2.32 |
| 8 | .alpha.-Methyl-.alpha.-[4-methyl-3-pentenyl]oxiranemethanol | 7.936 | 0.92 |
| 9 | 2-Carene | 8.165 | 0.96 |
| 10 | Linalool | 8.419 | 27.34 |

### Toxicity Study

Fumigation toxicity of EO was evaluated against adults at 40, 80, 120, 160 and 200 μL/L concentrations. At 160 μL/L of EO showed 100% of adult beetle’s mortality at 9 h of treatment. More than 91.56% of mortality recorded at 120 μL/L concentration during 24 h observation period. At the lowest concentration (40μL/L) of *C. citrinus* oil exhibited 50% mortality after 24 h of exposure, and there was a gradual increase in insect mortality while increasing concentrations of EO. The lethal concentration (LC_50_) of *C. citrinus* oil against *T. castaneum* adults after 24 h of exposure was 37.05 μL/L (Table 2a). The overall results, *C. citrinus* essential oil showed a time and concentration related effect against *T. castaneum*.

**Table 2a.** Fumigation toxicity (%) of essential oil from *C. citrinus* against *T. castaneum*.

| Concentrations (µL/L) | Period of study (hrs) |  |  |  |  |
| --- | --- | --- | --- | --- | --- |
|  | 3 | 6 | 9 | 12 | 24 |
| 40 | 0.0±0.0<sup>a</sup> | 12.0±2.00<sup>a</sup> | 30.0±3.16<sup>a</sup> | 36.0±2.45<sup>a</sup> | 50.0±1.75<sup>a</sup> |
| 80 | 0.0±0.0<sup>a</sup> | 16.0±2.45<sup>a</sup> | 58.0±2.00<sup>b</sup> | 68.0±3.74<sup>b</sup> | 81.3±1.93<sup>b</sup> |
| 120 | 0.0±0.0<sup>a</sup> | 30.0±3.16<sup>b</sup> | 72.0±3.74<sup>c</sup> | 82.0±2.00<sup>c</sup> | 91.6±2.12<sup>c</sup> |
| 160 | 8.0±3.74<sup>b</sup> | 48.0±3.74<sup>c</sup> | 100.0±0.00<sup>d</sup> | 100.0±0.00<sup>d</sup> | 100.0±0.00<sup>d</sup> |
| 200 | 22.0 ±2.00<sup>b</sup> | 64.0±2.45<sup>d</sup> | 100.0±0.00<sup>d</sup> | 100.0±0.00<sup>d</sup> | 100.0±0.00<sup>d</sup> |
| Shapiro-Wilk test | ns | ns | ns | ns | ns |

Mean of five replication ± SE

### Repellency - Larvae

Larvicidal activity of EO was studied at different concentrations *viz.,* 40, 80, 120, 160, 200, 240, 280 and 320 μL/L. Maximum larvicidal activity of 95.78% was observed at 320 μL/L concentration on 48 hrs after exposure period, while the lowest concentration (40 μL/L) exhibited 16.89% larvicidal activity. More than 50% larvicidal activity was recorded at 160 μL/L during 48 h after treatment. The lethal concentration (LC_50_) of *C. citrinus* oil against *T. castaneum* larvae was 144.31 μL/L for 48 h of exposure (Table 2b). The fumigation toxicity of *C. citrinus* EO was concentration and time dependent against *T. castaneum* larvae, however, the larvae appeared to be more tolerance to the EO than adults.

**Table 2b.** Larvicidal activity (%) of essential oil from *C. citrinus* against *T. castaneum*.

| Concentrations<br>($\mu\text{L/L}$) | Period of study (hrs) | | | | | | |
| --- | --- | --- | --- | --- | --- | --- | --- |
|  | 3 | 6 | 9 | 12 | 24 | 36 | 48 |
| 40 | 0.0±0.0 | 0.0±0.0<sup>a</sup> | 2.0±2.00<sup>a</sup> | 4.0±2.45<sup>a</sup> | 6.0±2.45<sup>a</sup> | 10.2±3.16<sup>a</sup> | 16.9±2.38<sup>a</sup> |
| 80 | 0.0±0.0 | 0.0±0.0<sup>a</sup> | 4.0±2.45<sup>ab</sup> | 14.0±2.45<sup>b</sup> | 18.2±3.62<sup>b</sup> | 22.9±1.84<sup>b</sup> | 31.6±3.99<sup>b</sup> |
| 120 | 0.0±0.0 | 0.0±0.0<sup>a</sup> | 6.0±2.45<sup>abc</sup> | 18.0±3.74<sup>bc</sup> | 22.4±1.93<sup>bc</sup> | 26.9±3.69<sup>b</sup> | 42.2±3.76<sup>bc</sup> |
| 160 | 0.0±0.0 | 6.0±2.45<sup>abc</sup> | 10.0±3.16<sup>abcd</sup> | 22.0±2.00<sup>bcd</sup> | 26.2±3.77<sup>bc</sup> | 31.1±2.87<sup>bc</sup> | 50.9±3.07<sup>c</sup> |
| 200 | 0.0±0.0 | 8.0±2.00<sup>bc</sup> | 14.0±2.45<sup>bcd</sup> | 26.0±2.45<sup>cde</sup> | 30.4±2.82<sup>bcd</sup> | 35.1±3.47<sup>bc</sup> | 55.3±3.85<sup>c</sup> |
| 240 | 0.0±0.0 | 12.0±2.00<sup>cd</sup> | 16.0±2.45<sup>cde</sup> | 30.0±3.16<sup>de</sup> | 34.7±2.26<sup>cd</sup> | 41.8±1.09<sup>cd</sup> | 70.4±3.39<sup>d</sup> |
| 280 | 0.0±0.0 | 14.0±2.45<sup>cd</sup> | 20.0±3.16<sup>de</sup> | 36.0±2.45<sup>ef</sup> | 40.7±2.66<sup>de</sup> | 47.8±3.34<sup>d</sup> | 80.9±2.06<sup>d</sup> |
| 320 | 0.0±0.0 | 18.0±3.74<sup>d</sup> | 26.0±2.45<sup>e</sup> | 42.0±2.00<sup>f</sup> | 49.1±2.51<sup>e</sup> | 64.7±2.00<sup>e</sup> | 95.8±2.59<sup>e</sup> |
| Shapiro wilk test | s | s | s | s | s | s | s |

Mean of five replication ± SE

**Table 2c.** Lethal concentration of essential oil from *C. citrinus* against *T. castaneum* adult and larvae.

| Stage of the insect | LC<sub>50</sub><br>(μL/L) | 95%Feducial level |  | LC<sub>90</sub><br>(μL/L) | 95%Feducial level |  | Chi-square |
| --- | --- | --- | --- | --- | --- | --- | --- |
|  |  | Lower | Upper |  | Lower | Upper |  |
|  | 24 hrs after treatment |  |  |  |  |  |  |
| Adult | 37.05 | 14.09 | 51.02 | 102.82 | 89.70 | 123.41 | 3.74 |
|  | 48 hrs after treatment |  |  |  |  |  |  |
| Larva | 144.31 | 123.32 | 162.85 | 321.62 | 289.44 | 369.08 | 11.01 |

### Repellency - adult

Repellent activity of four different concentrations (5, 10, 15 and 20 μL/L) of *C. citrinus* EO was evaluated against *T. castaneum* adults using a Y-arm olfactometer. A 100% adult repellent activity was observed at 20 μL concentrations after 24 h. The lower concentration of EO exhibited more than 31.1% repellent activity (Table 3).

**Table 3:**
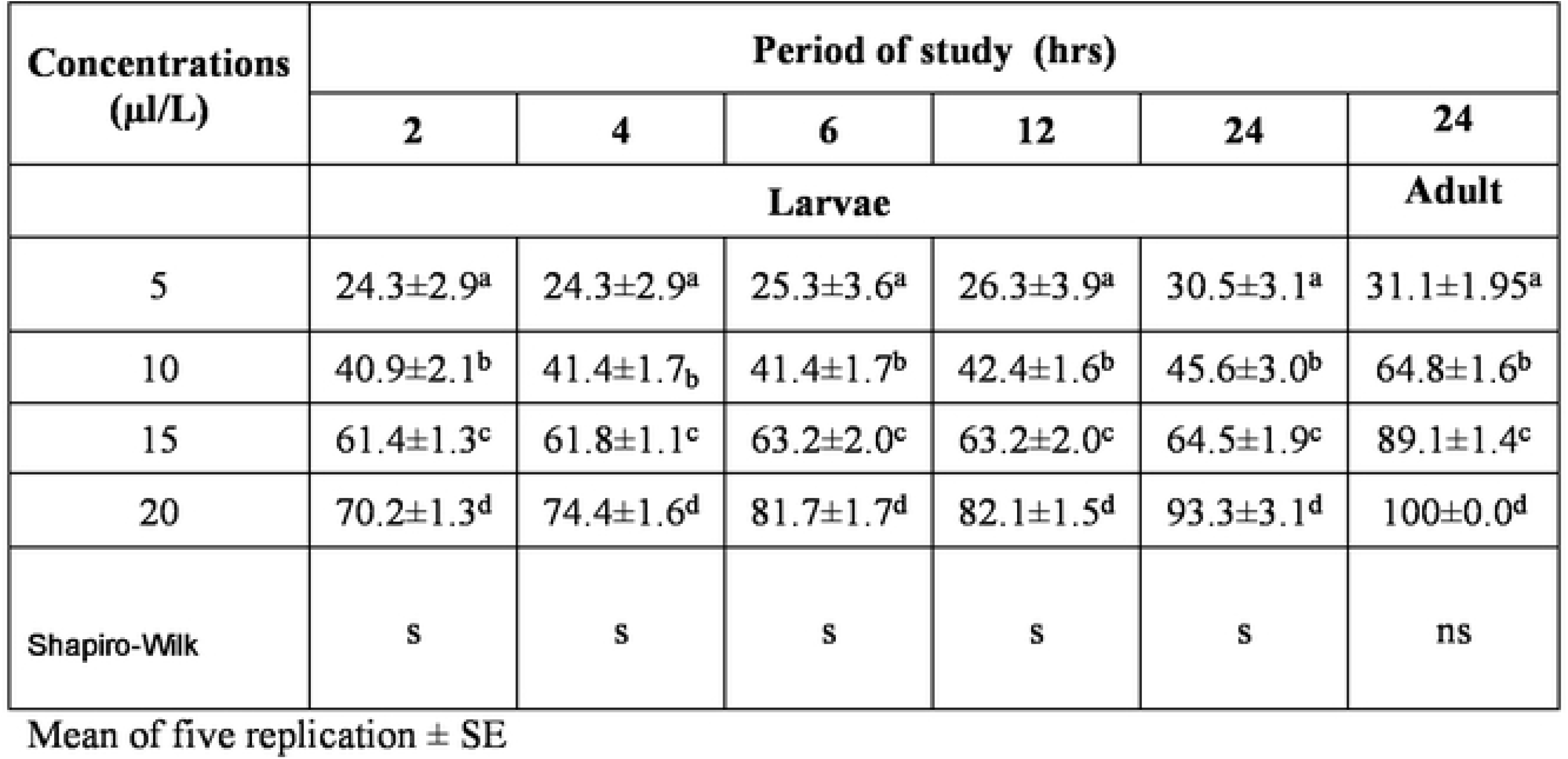
Repellent activity (%) of essential oil from *C. citrinus* against *T. castaneum*.

The larval repellency was conducted in Petri dishes using a choice-based method at 5, 10, 15 and 20 μL concentrations and different observation period of 2, 4, 6, 12, and 24 h. The maximum repellency (93.3%) was observed at 20 μL concentrations at 24 h observation. Lowest concentration showed 30% repellent activity against *T. castaneum* larvae at 24 h (Table 3). Overall, the results indicate that the EO exhibited good repellence potential on both larvae and adults.

### Fecundity and knock-down activity

The oviposition study was carried out at 5, 10, 20 & 30 μL/L concentrations of EO by fumigation toxicity. Fecundity in the control beetle group laid on an average of 5.8 eggs per individual. The concentration of 20 & 30 μL/L showed 2.6 and 1.4 eggs. In terms of per cent reduction; 20 & 30 μL/L concentrations showed 55.17 & 75.86% reduction in fecundity, respectively. *C. citrinus* EO significantly reduced oviposition deterrence activity at 20 & 30 μL/L (Table 4). Knockdown effect was increased according the increasing concentration. Maximum knockdown activity of 35.5% was observed at 30μL/L.

**Table 4:**
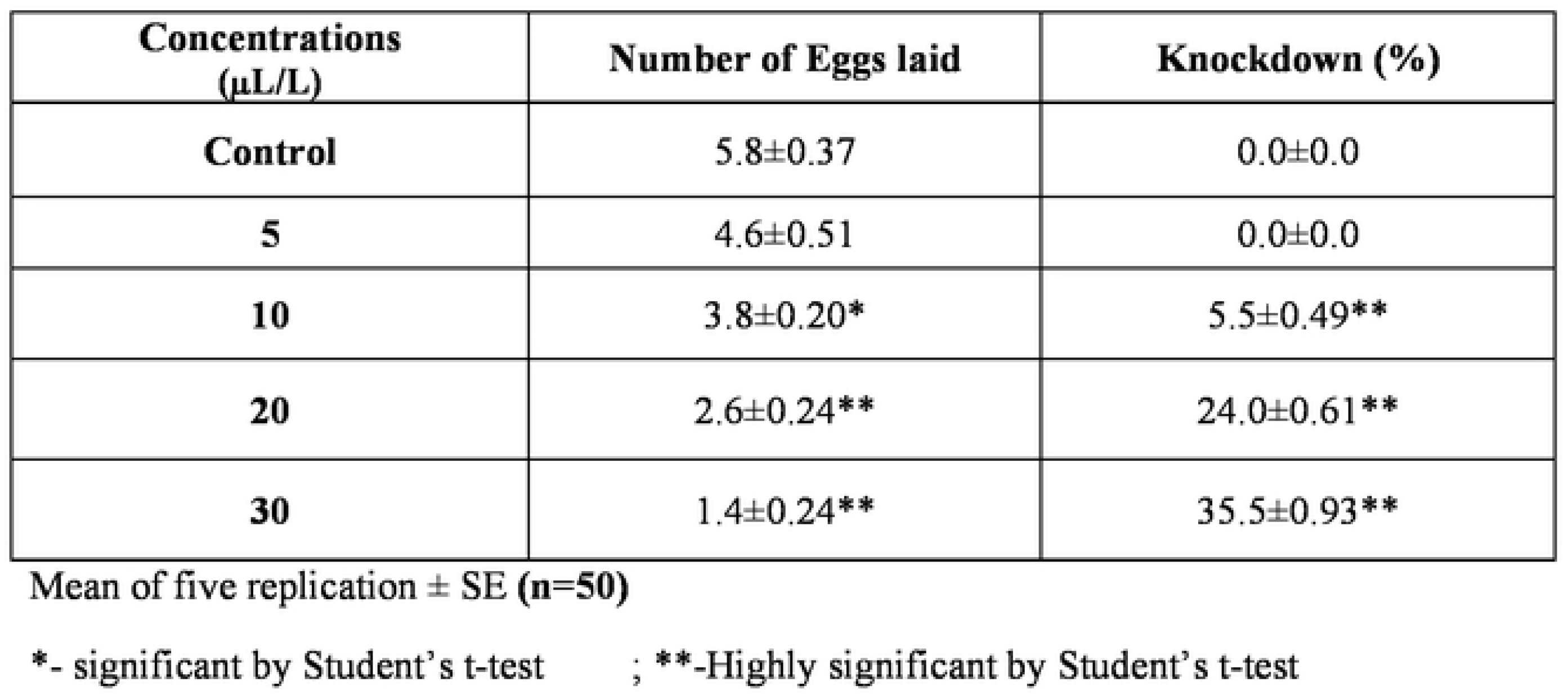
Oviposition deterrent (number of eggs/insect/day) of essential oil from *C. citrinus* on *T. castaneum*.

| **Concentrations**<br>**($\mu\text{L/L}$)** | **Number of Eggs laid** | **Knockdown (%)** |
| --- | --- | --- |
| **Control** | 5.8±0.37 | 0.0±0.0 |
| **5** | 4.6±0.51 | 0.0±0.0 |
| **10** | 3.8±0.20* | 5.5±0.49** |
| **20** | 2.6±0.24** | 24.0±0.61** |
| **30** | 1.4±0.24** | 35.5±0.93** |

Mean of five replication ± SE (n=50)
\*- significant by Student's t-test ; \*\*-Highly significant by Student's t-test

### Growth inhibition effects

#### Ovicidal activity and egg hatchability

The ovicidal activity of *C. citrinus*, EO was studied against *T. castaneum* at four different concentrations. Maximum ovicidal activity of 91.49% was observed at 30 μL/L concentration of EO.

The 5 μL/L concentration exhibited egg hatchability of 56.55% while control exhibited 89.15% egg hatchability. The minimum egg hatchability of 8.51% was recorded at 30 μL/L concentration (Table 5).

**Table 5:** Bioefficacy of essential oil from *C. citrinus* against different life stages of *T. Castaneum*.

| **Concentrations** | **Ovicidal activity (%)** | **Egg hatchability (%)** | **Larval survival (%)** | **Adult emergence of F1 generation (%)** |
| --- | --- | --- | --- | --- |
| **Control** | 10.85±0.25 | 89.15±0.25 | 86.96±1.23 | 80.58±1.15 |
| **5** | 43.45±0.23 | 56.55±0.24** | 70.67±0.57* | 53.45±0.27** |
| **10** | 56.78±0.36 | 43.29±0.36** | 60.61±1.41** | 36.14±0.90** |
| **20** | 78.69±0.15 | 21.31±0.15** | 51.04±1.22** | 21.82±0.67** |
| **30** | 91.49±0.13 | 8.51±0.13** | 42.54±1.32** | 5.15±2.10** |

Mean of five replication ± SE;\*- significant by Student's t-test; \*\*-Highly significant by Student's t-test

### Larvae survival and adult emergence

Larval survival, and adult emergence (F_1_ generation) were 86.96% and 80.58%, respectively in the control group. In contrast, the 30 μL/L concentration of EO exhibited larval survival 42.54%, and it was notably lower than the control. The 30 μL/L treatment allows 5.15% of adult to emerge, when compared to control concentration A significant reduction in adult emergence 5.15% was observed at 30 μL/L.

### Quantitative analysis of *T. castaneum* adult biochemistry

Based on the obtained results, sub-lethal concentrations (5, 10 and 20 μL/L) were used to study the impact of *C. citrinus* EO on various biochemical constituents in adult *T. castaneum* beetles. Results indicated that the biochemical constituents measured in *T. castaneum* adult beetles can significantly vary after exposure to sub-lethal concentrations of EO for 24 h. The total protein content of *T. castaneum* adult was highly and significantly reduced relative to the control value of 7.43 mg**/**mL of protein to 6.19 mg**/**mL, 5.72 mg**/**mL, 5.32 mg**/**mL in adults exposed to different concentrations (5, 10 and 20 μL/L) of EO, respectively (Fig. 1a). Acetylcholinesterase activity dramatically increased in the 10 and 20 μL/L, treatments when compared to the control; while, at 20 μL/L concentration treatment reduced Acetylcholinesterase activity (Fig. 1b).

**Fig 1a.**
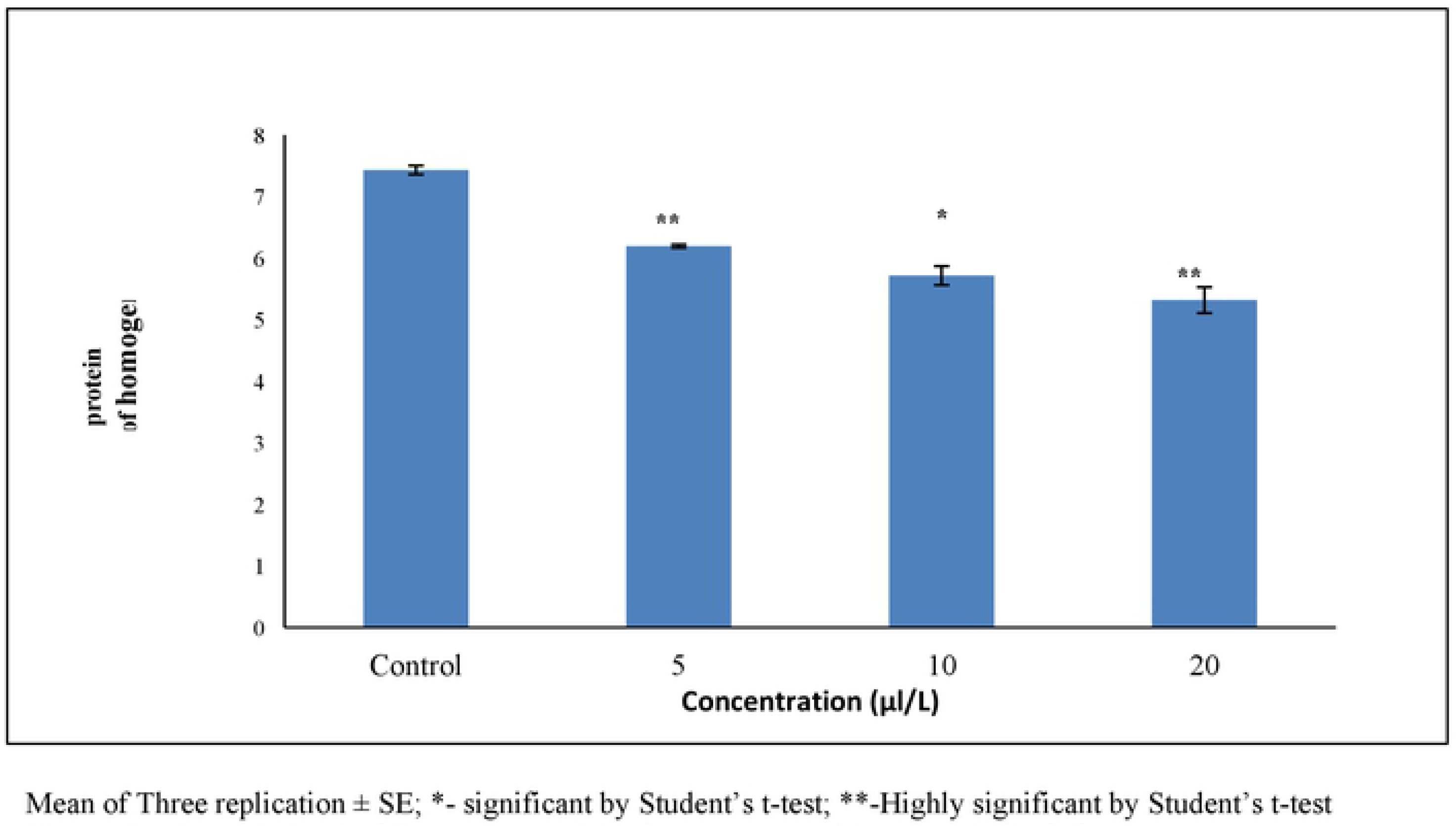
Total protein level of *T. castaneum* adult after treatment with essential oil of *C. citrinus*

**Fig 1b.**
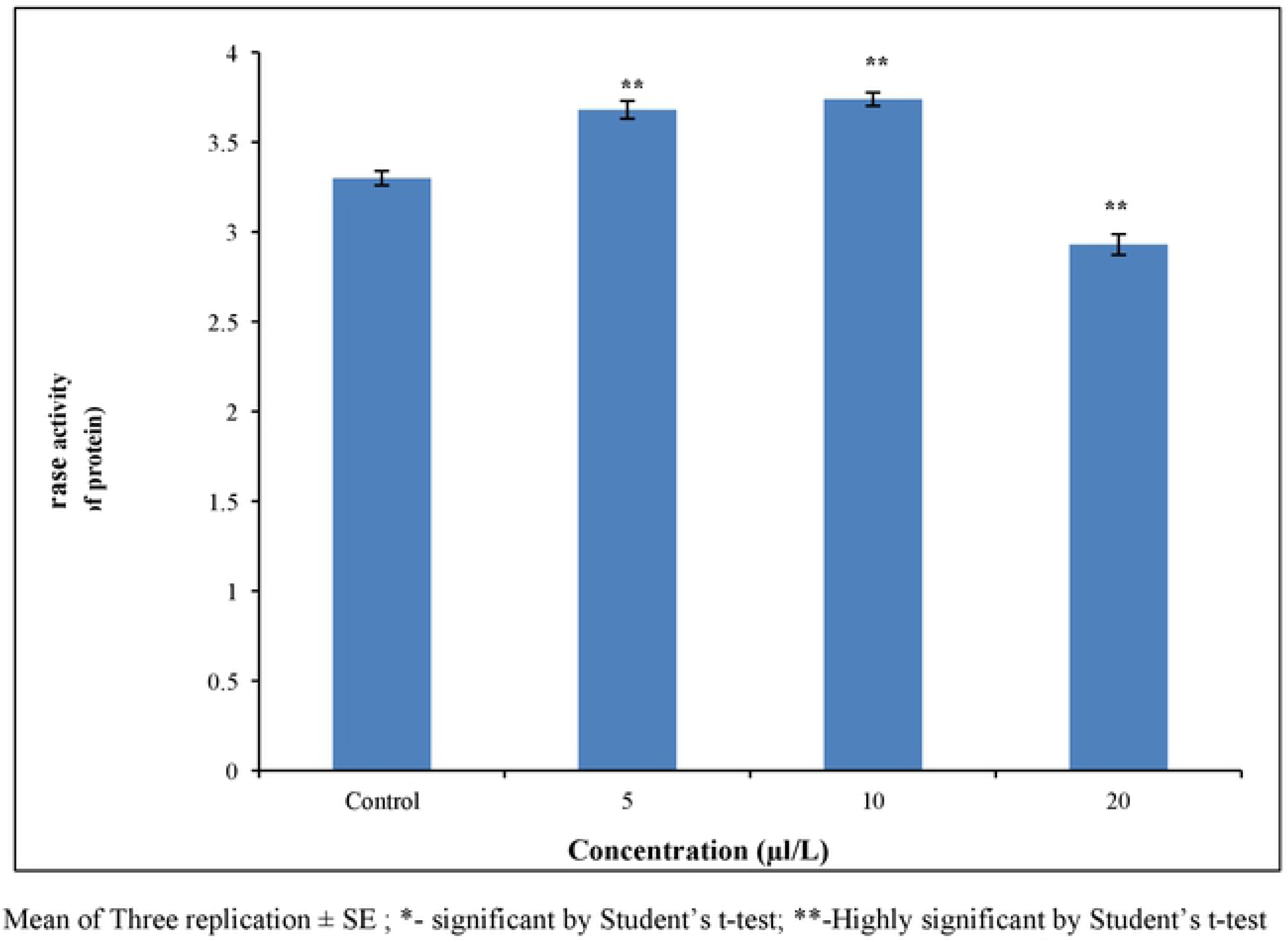
Acetylcholinesterase activity of adult (mM ACT released/min/mg protein) *T. castaneum* adult after treatment with essential oil of *C.citrinus*

The level of α-Carboxylesterase activity was significantly increased at three of the selected sub-lethal concentrations of EO, compared to the control (Fig. 1c). β-carboxylesterase activity level was also drastically elevated in the 5, 10 and 20 μL/L treatments compared to the control, however, no significant difference was observed between the different sub-lethal treatments (Fig. 1d).

**Fig 1c.**
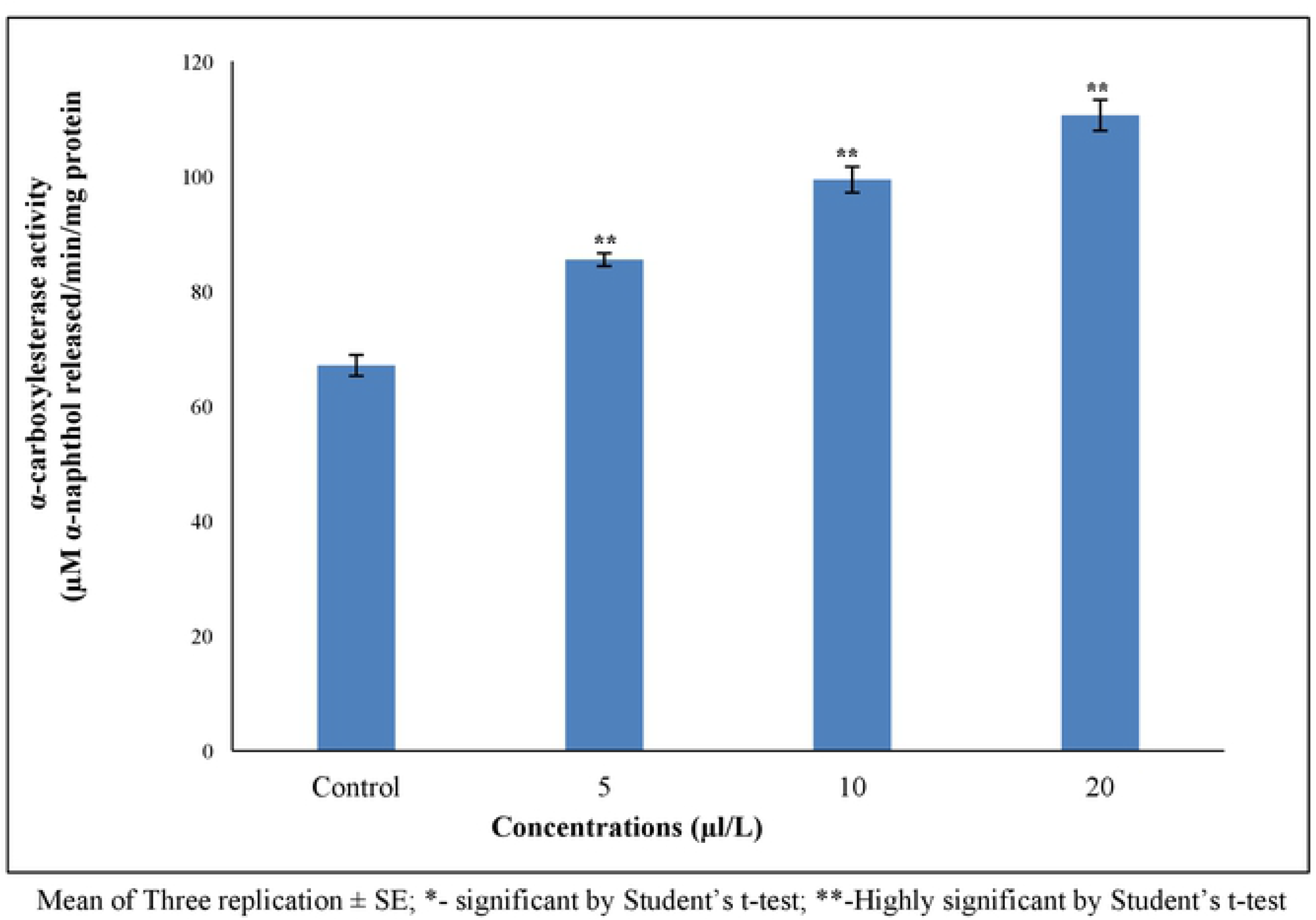
α-Carboxylesterase activity (μM α-naphthol released/min/mg protein) of *T. castaneum* adult after treatment with essential oil of *C. citrinus*

**Fig 1d.**
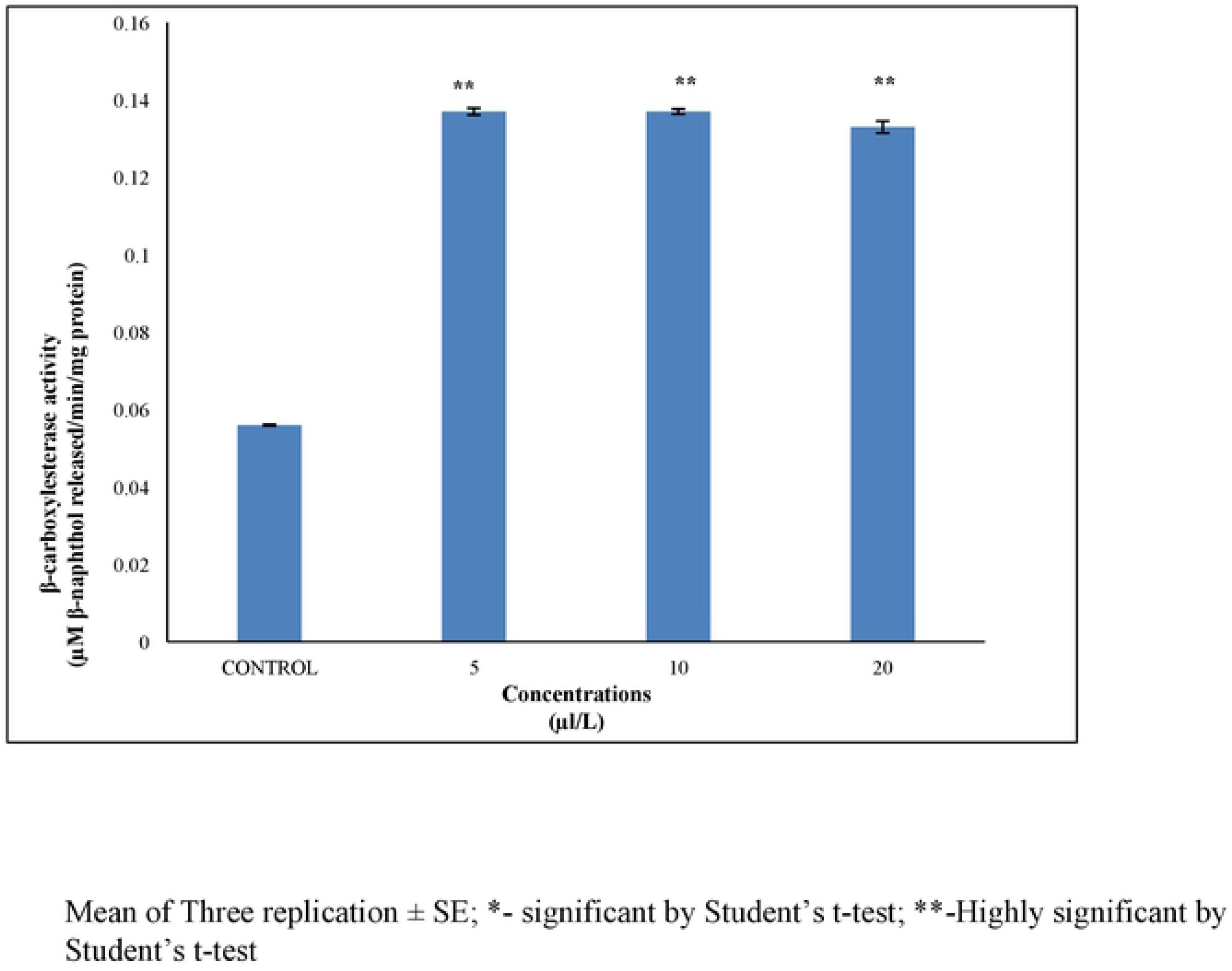
β-Carboxylesterase activity (μM β-naphthol released/min/mg protein) of *T. castaneum* adult after treatment with essential oil of *C. citrinus*

Exposure of *T. castaneum* adults to *C. citrinus* EO resulted in decreased level of acid phosphatase at selected concentrations when compared to control group (Fig. 1e). Alkaline phosphatase activity was significantly reduced in the 5 μL/L treatment and drastically elevated in the 10 μL/L treatment. Significantly lower activity was recorded in the 20 treatment when compared to the control group (Fig. 1f). Glutathione-S-Transferase levels significantly increased in the 10 and 20 μL/L treatments, relative to the control group, but was significantly lower, relative to the control, in the 10 μL/L treatment (Fig. 1g).

**Fig 1e.**
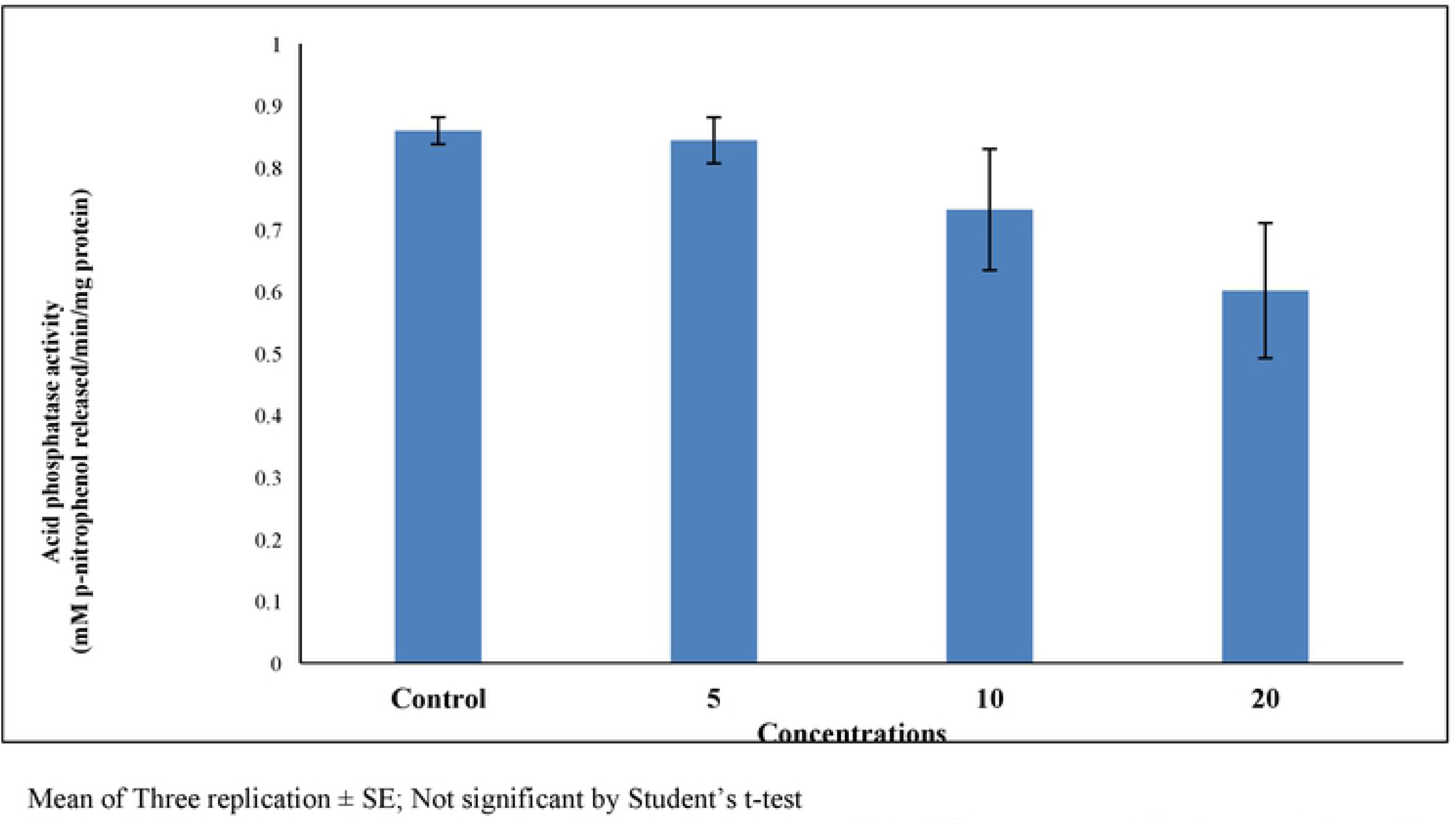
Acid phosphatase (mM p-nitrophenol released/min/mg protein) of *T. castaneum* adult after treatment with essential oil of *C. citrinus*

**Fig 1f.**
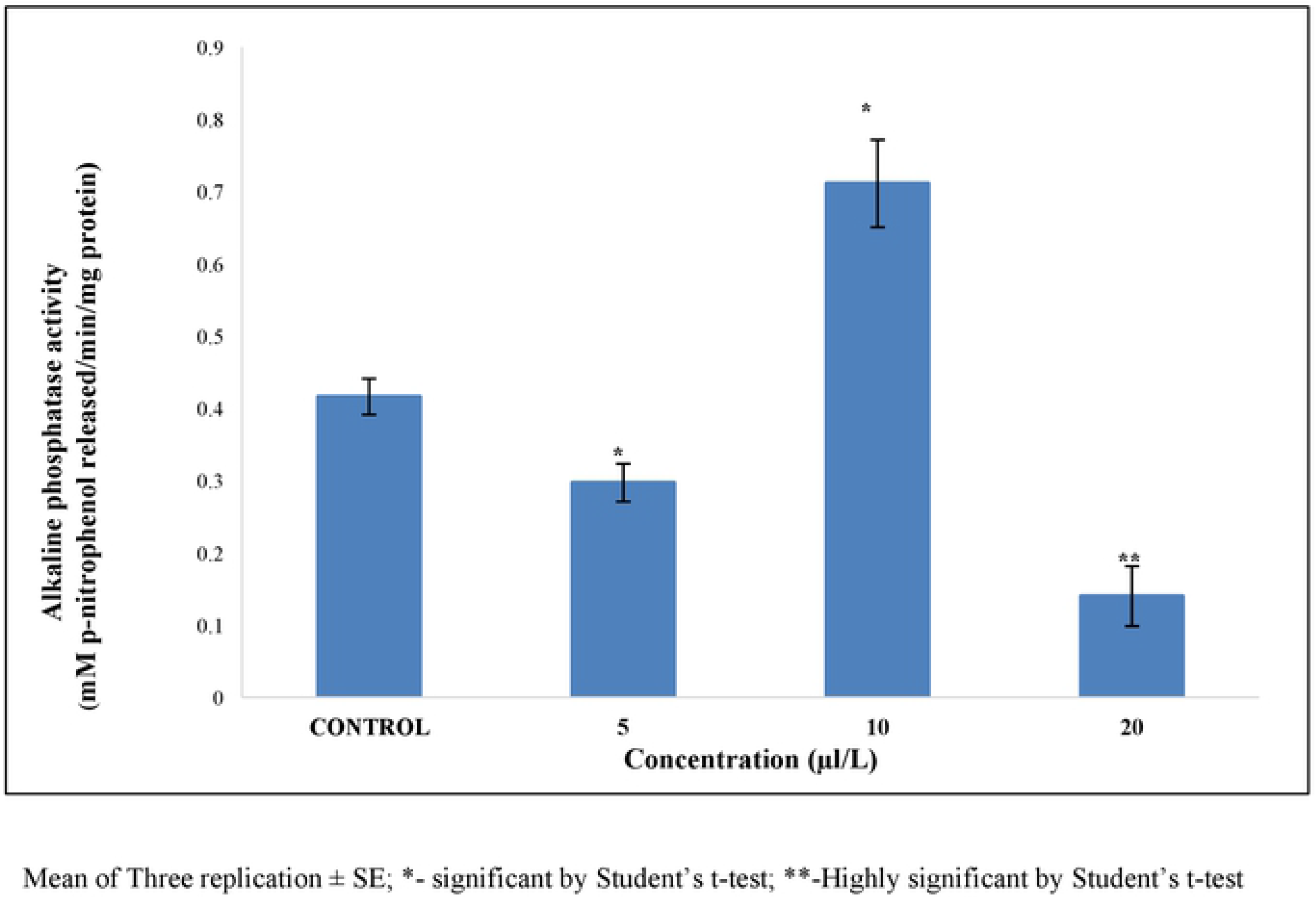
Alkaline phosphatase(mM p-nitrophenol released/min/mg protein) of *T. castaneum* adult after treatment with essential oil of *C. citrinus*

**Fig 1g.**
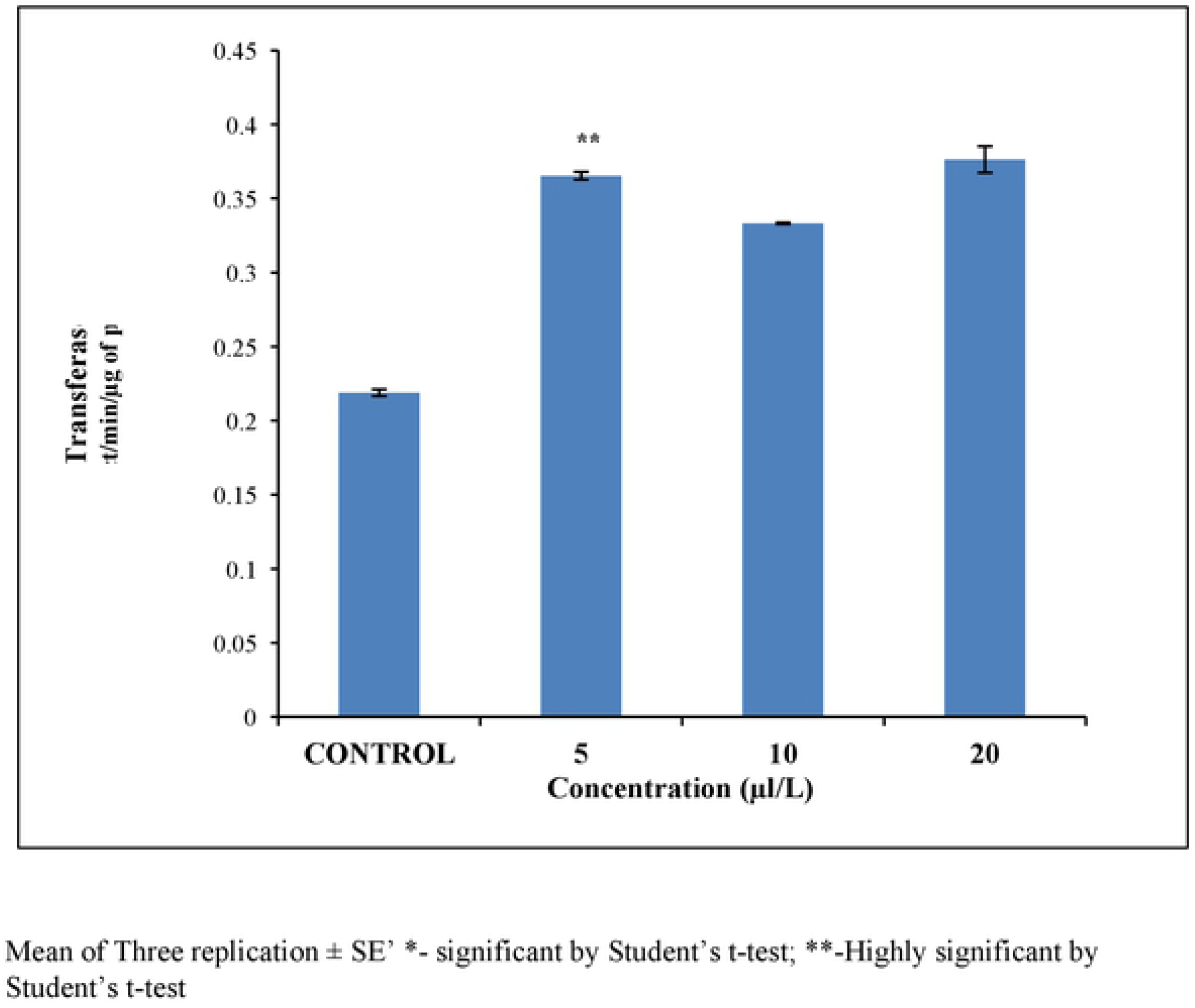
Glutathione-S-Transferase activity (CDNB product/min/mg protein) of *T. castaneum* adult after treatment with essential oil of *C. citrinus*

### 3.9 Qualitative analysis of T. castaneum adult biochemistry

A qualitative analysis of total proteins was analysed using the native PAGE method. Protein extracted from *T. castaneum* adults treated with sub-lethal concentrations of *C. citrinus* EO showed a reduction in the number protein bands, relative to the control (Fig. 2a). The intensity of the esterase band of β-Carboxylesterase isoenzyme was modulated by the concentration of EO. The intensity of the band was lowered in the 10 treatment but increased gradually as the concentration of EO increased. The two lower isoenzyme bands decreased in their intensity, relative to the control, at the lower concentrations of EO and gradually increased when beetles were exposed to higher concentration of EO (Fig. 2b). Electrophoretic analysis of acid and alkaline phosphatase enzyme activity in adult beetles was not affected by exposure to the range of sub-lethal concentrations of EO used in the experiment (Fig. 2c,d).

**Fig 2a.**
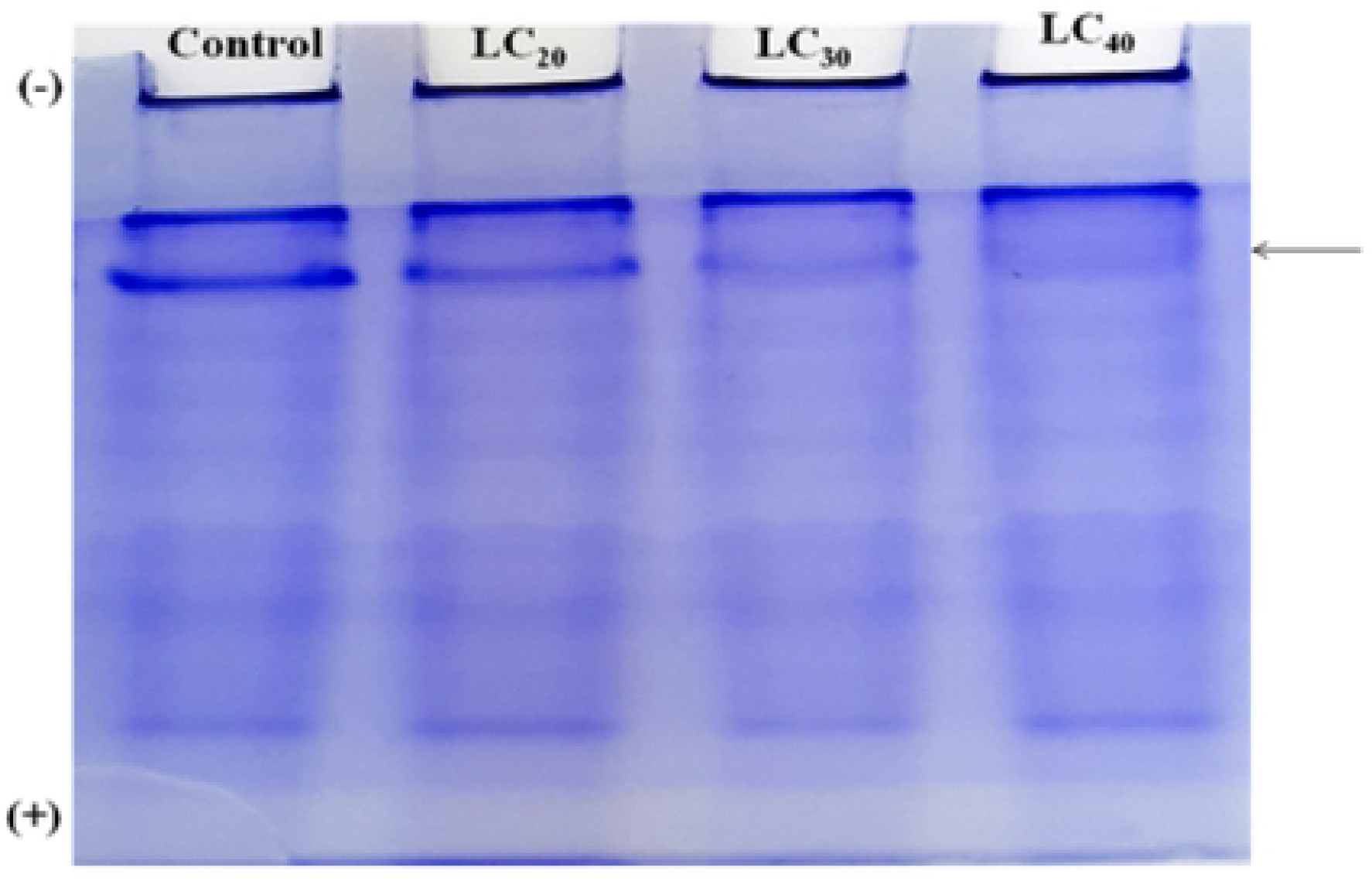
Qualitative analysis of total protein in native PAGE, of *T. castaneum* adult after treatment with essential oil of *C. citrinus*

**Fig 2b.**
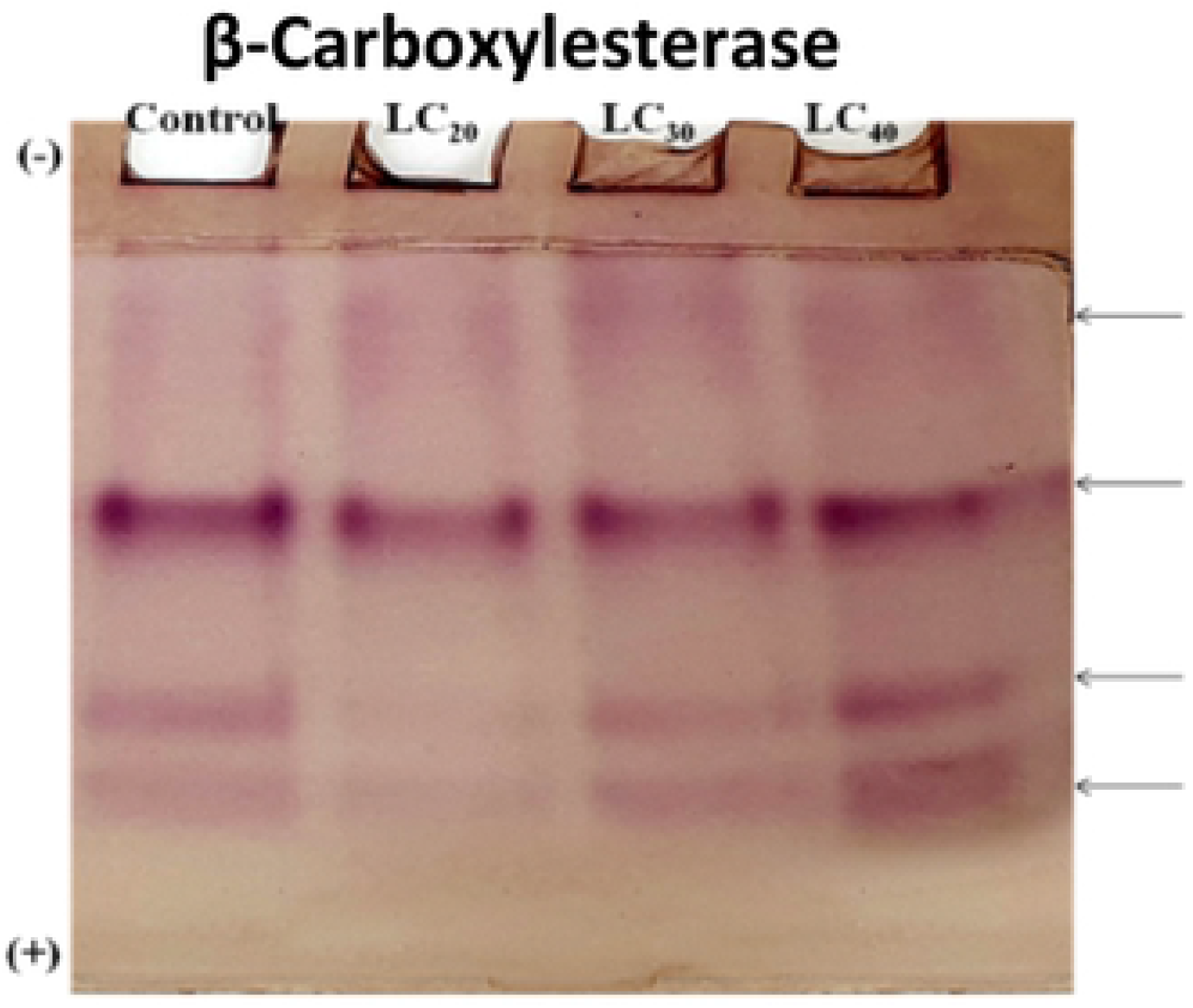
Qualitative analysis of isoenzyme of β-Carboxylesterases of *T. castaneum* adult after treatment with essential oil of *C. citrinus*

**Fig 2c.**
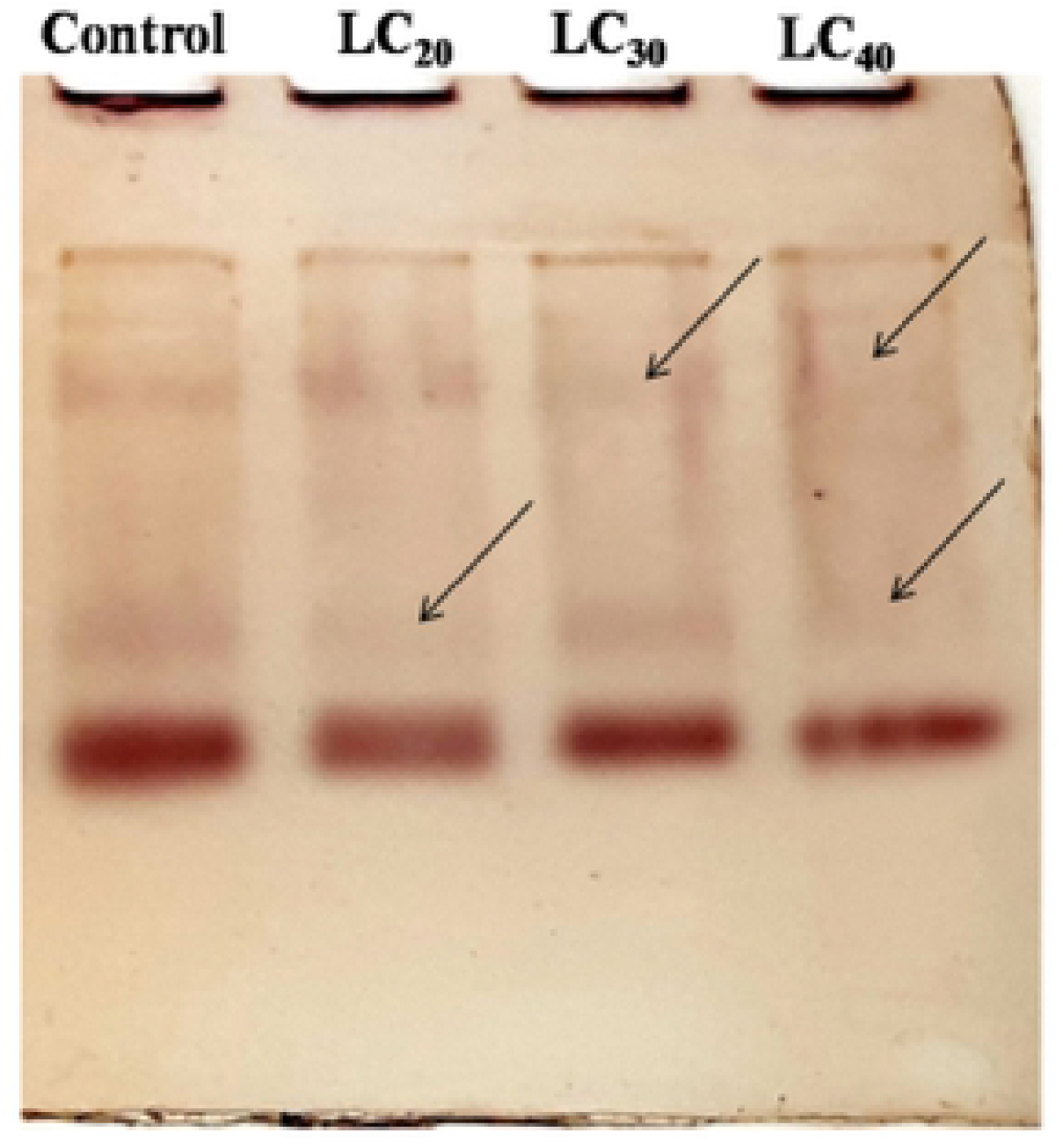
Qualitative analysis of acid phosphatises of *T. castaneum* adult after treatment with essential oil of *C. citrinus*

**Fig 2d.**
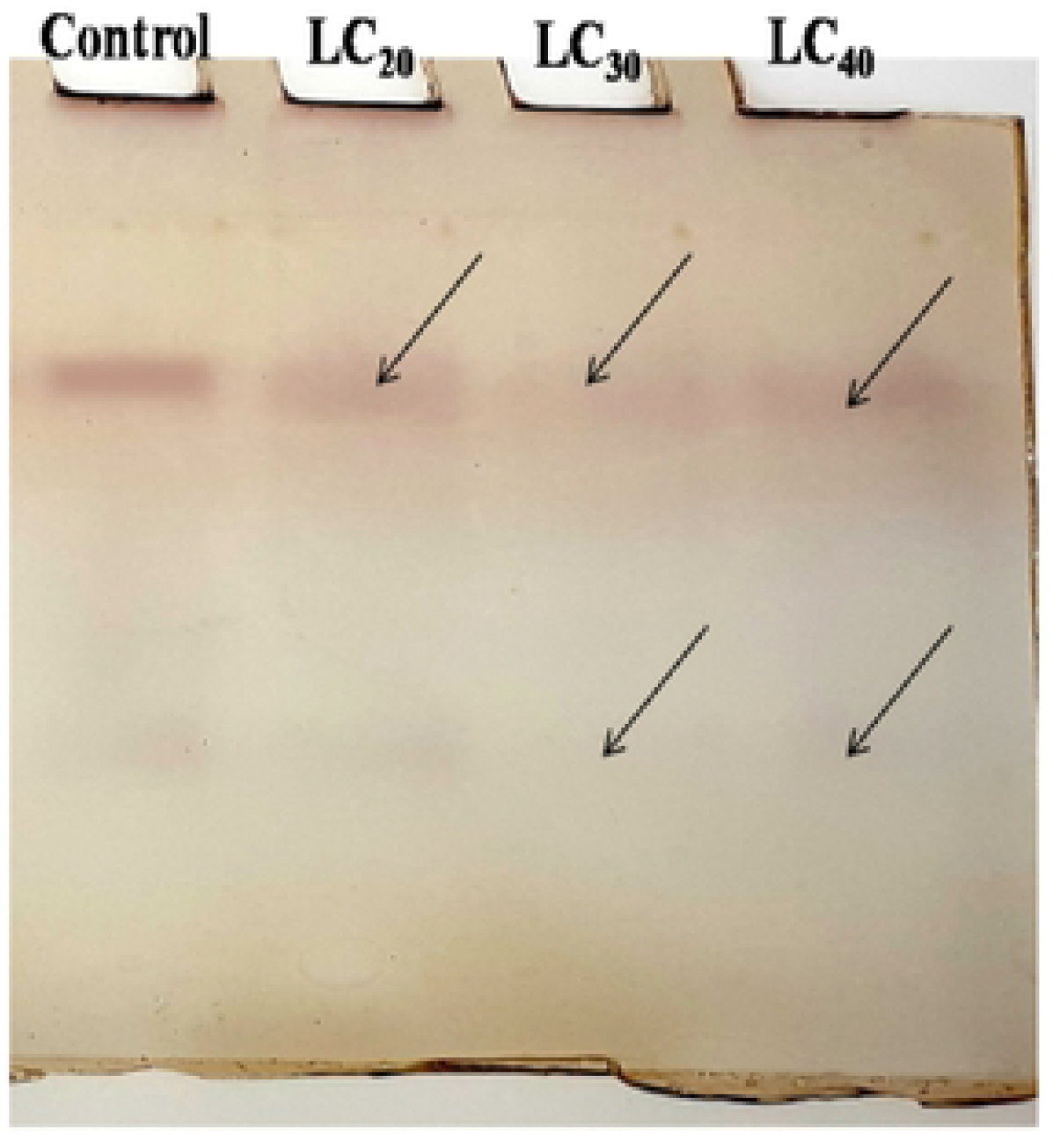
Qualitative analysis of alkaline phosphatases *T. castaneum* adult after treatment with essential oil of *C. citrinus*

## Discussion

Botanical pesticides have the potential to eradicate pests without causing harm to the environment and non-target organisms. Since botanicals are generally biodegradable and do not leave persistent residue, they have been increasingly used in recent years. Essential oils obtained from plants have been shown to be very effective on insect pests, especially in stored grains. Since they are generally volatile in nature, essential oils are very easy to use and kill the pests of stored grains. In the present study, essential oil extracted from *C. citrinus* leaves was tested for its potential against the larvae and adult beetles of *T. castaneum*.

GC-MS analysis of *C. citrinus* revealed six different compounds, in which eucalyptol was the major constituent (40.44%), followed linalool (27.34%), and alfa- Pinene (17.46%). However, GC-MS analysis of EO from the same plant from Ethiopia revealed 15 different compounds, which also included eucalyptol as the major constituent (76.9%) [37]. In contrast, EO extracted from the same plant growing in Western Himalayas contained only 9.8% eucalyptol [38]. These data clearly indicated that the level and type of constituents in EO extracted from this plant varies depending on where the plant was collected, and most likely the climate and ecology of that region. These results are in accordance with Misharina [39], Souza and Vendramim [40], Isikber et al. [41],who indicated that the variations in extracted compound’s, could be associated with geographical location, collection time, amount of sunlight, length of storage, temperature, and extraction methods.

Fumigation is one of the most effective, practically feasible, and rapid methods that can be used to protect feedstuffs, stored food grains and other agricultural products from pest infestation [42–43]. Many plant essential oils and their constituent compounds have been reported to have fumigant activity. Essential oils of *Artemisia annua* [44]*, Lipia alba* [45], *Curcuma longa* [46], *Cinnamomum camphora, Eucalyptus globules* [47], *Boswellia carterii* [48], *C. camphora, Myristica fragrans, Rosmarinus officinalis* [11] and *Mentha piperita* [13], are reported to have biocidal activity against stored grain pests. Bioactivity can vary greatly, however, due to the variability in chemical composition of each EO and the stage of plant growth and plant organ that was used for extraction. In the present study, fumigation with EO extracted from leaves of *C. citrinus* resulted in 100% mortality in adult beetles of *T. castaneum* at 9 hrs after treatment at a concentration of 160 μL/L. Similar results were obtained from the use of *Coriandrum sativum* seed oil, where mortality in *C. maculatus* and *T. confusum* increased with the use of increasing concentrations of EO from 43 to 357 *μ*L/L air [49]. The EO used in our study exhibited different levels of toxicity to larvae vs. adult beetles. In general, fumigation toxicity was lower against larvae than adult beetles. Earlier, Liu and Ho [50]; Huang et al. [51], Tripathi et al.[46]; Isikber et al. [41] have been reported similar results.

Several plant essential oils exhibit feeding deterrence, acute toxicity, repellency and developmental disruption in many storage insect pests due to the complex combinations of monoterpenoids and allied phenolic compounds present in essential oils [52–55]. β-pinene has been reported to exhibit strong toxicity and repellent activity against *T. castaneum* adults [56]. α-cymene, α-terpinene, α-terpeneol, and terpine-4-ol exhibited fumigant activity against *S. oryzae* [57]. In general, the major volatile components in an essential oil are responsible for its activity. As demonstrated by Maciel et al. [58], eucalyptus oil, in which1, 8-cineole is the major component, exhibited strong biological activity. Our results are in accordance with earlier literature where e it stated that eucalyptol, linalool, and terpine-4-ol are present as major constituents of the selected EO. Thus, a very high degree of fumigation, repellent, and other biological activity was observed.

The number of eggs laid by adult beetle was significantly (P < 0.05) lowered by exposure of adult beetles to cardamom oil at 16 and 21 mg/cm^2^ concentrations [59]. In the present study, a significant reduction in oviposition was recorded when adult beetles were exposed to different concentrations of EO. Similarly, Moura et al. [22] also reported that EO derived from *Vanillosn opsis arborea* reduced the level of oviposition when compared to control. A previous report indicated the decreased oviposition (28%) at 5.2 mg/cm^2^ concentration when the adult beetles were exposed to EO derived from leaves of *C. loga*. The reduction in oviposition was probably due to physically weakened insects as well as lesser surviving insects [46].

The inhibition of egg hatchability or ovicidal activity was rapidly exhibited by exposure to the EO without any direct contact with the eggs due the volatiles released by the EO. The vapour of essential oils has been shown to diffuse through the permeable membranes of insect eggs into the chorion and vitelline membrane [60–61]. The diffusion of essential oil vapours into the eggs results in a disruption in normal physiological and biochemical processes [62].The ovicidal activity observed in the present study confirms the above statements, as the tested EO resulted, 91.49% ovicidal activity in *T. castaneum*.

The suppression and reduction of the F1 generation could be due to the toxicity of EO to all of the all life stages of the insect, from eggs to adults, via both fumigant and possibly stomach action [59]. In the present study, a drastic reduction (94.85%) was observed in adult emergence when the eggs were exposed to EO.

Proteins are the most abundant organic compounds in the insect body as they provide structure and muscle to the insect body, transport substances into the haemolymph, provide energy, and catalyse chemical reactions in the form of enzymes [63–64]. Decreasing protein content is commonly occurs when the insects treated with lethal compounds [65]. Protein synthesis may be reduced or inhibited in response to prolonged toxic stress [66–68]. Insects degrade the protein content into amino acid and the release energy to compensate the lowering energy level during stress condition by Nath et al. [69]. Reductions in protein levels were observed in the present study and has been previously reported that protein content was reduced due to the toxicity of plant product [70–72,25].

Esterases are synthesized in insects during various development stages. The level of esterase activity is not constant throughout the life cycle. In the present study, AChE activity was inhibited by the higher concentrations of *C. citrinus* oil. Saponins are able to inhibit AChE and the inhibition increases with increasing concentration [73]. Inhibiting AChE results in the accumulation of acetylcholine at cholinergic synapses and causes hyper excitation of cholinergic pathways [74].

Carboxylesterase activity can be altered by plant secondary metabolites. Phenolic glycoside significantly increased the level of Carboxylesterase in *Lymantria dispar* [75]. A higher level of CarE activity was recorded in *Sitobion avenae* fed on diet with high in indole alkaloid content [76]. In the present study, both α- CarE and β- CarE levels increased in adult beetles exposed to *C. citrinus* EO.

Hydrolytic cleavage of phosphoric acid esters is catalysed by Phosphatase enzymes that are classified into “acid” or “alkaline” phosphatases based on their pH [77]. Acid phosphatases are alysosomal marker enzyme whose active site is in the gut of insects [78–82]. Alkaline phosphatases are a brush border membrane marker [83]. Exposure to plant compounds reduced the acid and alkaline phosphatises content in *Cnaphalocrocis medinalis* larvae [84]. A methanolic extract of *Melia azedarach* reduced the acid and alkaline phosphatises in fourth instar larvae of *C. medinalis* [85]. Similarly, *C. citrinus* EO also reduced both acid and alkaline phosphatase activity in a concentration dependent manner.

Glutathione transferases are the enzymes that catalyse the detoxification of insecticides typically after the phase-I metabolic process [86]. In the present study, elevated GST levels were observed in adult beetles exposed to the higher concentrations of *C. citrinus* EO. Shojaei et al [87] reported that adults of *Tribolium castaneum* exposure to *Artemisia dracunculus* EO enhanced the level of Glutathione-S-Transferase in a concentration dependent manner.

## 5. Conclusion

Essential oils exhibits wide range of biological activities which includes fumigant, repellent, oviposition and growth inhibitory activity, and act up on all insect development stages. Therefore, the potential of resistance development is very low. The present study provided promising results on the use of an EO extracted from leaves of *C. citrinus* against all life stages of the beetle, *T. castaneum*. Importantly, this potent essential oil may be useful for controlling beetle infestations in stored grains. However, further research will be required to address safety concerns regarding its effect on human health and the environment, as well to develop suitable formulations that improve insecticidal activity and reduce production costs.

## Acknowledgement

The present study was funded by the DST (DST-PURSE Phase II Programme), India, New Delhi sanctioned to Dr. M. Jayakumar Assistant Professor University of Madras. The authors would like to extend their sincere appreciation to the Deanship of Scientific Research at King Saud University for funding this research group NO (RG-1435-014).

